# Mesoscopic Oblique Plane Microscopy via Light-sheet Mirroring

**DOI:** 10.1101/2023.08.10.552834

**Authors:** Stephan Daetwyler, Bo-Jui Chang, Bingying Chen, Felix Zhou, Reto Fiolka

## Abstract

Understanding the intricate interplay and inter-connectivity of biological processes across an entire organism is important in various fields of biology, including cardiovascular research, neuroscience, and developmental biology. Here, we present a mesoscopic oblique plane microscope (OPM) that enables whole organism imaging with high speed and subcellular resolution. A microprism underneath the sample enhances the axial resolution and optical sectioning through total internal reflection of the light-sheet. Through rapid refocusing of the light-sheet, the imaging depth is extended up to threefold while keeping the axial resolution constant. Using low magnification objectives with a large field of view, we realize mesoscopic imaging over a volume of 3.7×1.5×1 mm^3^ with ∼2.3 microns lateral and ∼9.2 microns axial resolution. Applying the mesoscopic OPM, we demonstrate *in vivo* and *in toto* whole organism imaging of the zebrafish vasculature and its endothelial nuclei, and blood flow dynamics at 12 Hz acquisition rate, resulting in a quantitative map of blood flow across the entire organism.

## 1. Introduction

Mesoscopic fluorescence 3D imaging of model organisms in their entirety has emerged as an important imaging application in the life sciences. This paves the way to monitor systemic properties such as blood flow, every neuron in a behaving organism, or observe the metastatic cascade longitudinally in xenograft models. Moreover, advances in tissue clearing [1] and expansion [2] result in ever growing sample volumes that are rendered optically transparent. These applications require efficient 3D microscopes that possess rapid yet gentle volumetric acquisition capabilities and have sufficient spatial resolution, volumetric coverage, and optical sectioning capability.

Light-sheet fluorescence microscopy (LSFM) [3, 4] has the right attributes for the aforementioned tasks, as it offers parallelized and efficient 3D imaging combined with intrinsic optical sectioning, and in recent implementations also provides high volumetric acquisition rates [5-7]. Normally, LSFM uses separate optics for illumination and fluorescence detection, which can complicate sample access, especially for large samples or high-throughput applications. As an alternative, open-top light-sheet geometries are gaining popularity [8-12], as they leave in principle “infinite” lateral accessibility for large tissue slices, or series of organoids or model organisms in multi-well plates. Among open top architectures, oblique plane microscopy (OPM) is attractive as it employs only one primary objective for light-sheet illumination and fluorescence detection [13]. As such, the optical axis of the objective can be orthogonal to a coverslip or sample plate, a geometry for which objectives are usually designed and corrected for. In contrast, dual objective open-top LSFM architectures need aberration correction when interfacing with a coverslip at an off angle [12, 14]. In addition, OPM offers rapid optical scanning of volumes using galvo mirrors [5, 6], a speed potential that is especially attractive for functional imaging in the cardiovascular sciences, neurosciences, or when high throughput volumetric imaging is needed.

The concept of OPM, however, requires high numerical aperture objectives [13], as a large half opening angle is needed to give the light-sheets sufficient tilt, and to be able to capture fluorescence light with the downstream optical train. The latter constraint is rooted in how the tilted light-sheet plane is imaged onto a camera in OPM: Leveraging the principle of remote focusing [15], a distortion free and diffraction limited 3D image of the sample space is created by a secondary objective. When properly implemented, remote focusing maintains angles from the sample to the remote space, hence the fluorescence image of the light-sheet plane is tilted by the same angle as the light sheet emerges in sample space. A tertiary imaging system, whose focal plane overlaps with the remote image of the light-sheet plane, is then used to map the fluorescence onto a camera. Since the tertiary objective is tilted to the optical axis of the secondary objective, a light loss will occur, as the acceptance cones of the two objectives do not fully overlap. When using low NA objectives with a half opening angle below 30 degrees, this light-loss becomes total [16, 17]. In principle, one can increase the numerical aperture, and hence the acceptance angle of the tertiary objective to counter the light loss. However, there is an inverse tradeoff between field of view and numerical aperture [18], hence this approach is not well suited for mesoscopic imaging. Ideally, all three objectives should possess similar field of views to maximize volumetric coverage, and consequently all would have a similar numerical aperture.

Methods have been reported to “bend” light into the shallow acceptance cone of a tertiary objective in a mesoscopic OPM, such as reflective gratings [17], fiber face plates [16] or intentionally distorting the remote image space to make the image of the light-sheet less inclined [19]. While they provide solutions for low NA OPM implementations of the detection arm, they all come with specific caveats. Gratings with fine line spacings compress the light-cone in one spatial dimension, potentially lowering the resolution. In addition, higher diffraction orders may enter the tertiary objective in high numerical aperture scenarios. Fiber faceplates impose micron-sized resolution limits based on their fiber spacing. Lastly, compressing the remote space introduces spherical aberrations, which will lower the systems resolution.

Additionally, a low NA primary objective launches the light-sheet at a steep angle in sample space. This lowers the optical sectioning ability and axial resolution, as the light-sheet is far from being orthogonal to the optical axis of the detection objective as in a classical LSFM architecture. Augmenting the light-sheet tilt angle has been achieved using a secondary illumination objective [8, 20], or by placing a grating in front of the sample plane [19]. The former increases the complexity and alignment constraints, as two objectives now need to interface with the sample, negating the geometrical advantages of OPM over traditional LSFM. The latter comes with potential ghost images from higher diffraction orders, light-losses in the excitation and detection path, and a color dependent tilt angle of the light-sheet. As such, we found that there is still room for improvement for mesoscopic OPM designs.

Here, we introduce a reflection-based method to augment the light-sheet tilt angle. This is inspired by single objective selective plane illumination microscopy (soSPIM) [21], where a micro-mirror reflects the light-sheet by 90 degrees. We transfer this concept to mesoscopic OPM, where we increase the light-sheet tilt angle well beyond the half opening angle of the primary objective using a custom micro-prism. On the detection side, we leverage the concept of diffractive OPM (dOPM) to direct fluorescence light via a transmission grating to the tertiary objective. This resulted in a nearly isotropic lateral point spread function (PSF), as the grating overcomes “light-cone clipping” of traditional OPM systems and a low linespacing of the transmission grating leads to less light cone compression as in prior work in dOPM [17].

To further simplify the design and ease the alignment burden, we introduce a lens-less scanning system which dispenses with two scan lenses of a traditional OPM optical train. This makes our system more compact and easier to align, and we discuss the geometrical effects of this scanning mechanism in analytical detail.

Lastly we explore how we can increase the volumetric coverage in mesoscopic OPM through optical tiling while keeping the axial resolution constant [22]. To our knowledge, this is the first time this has been implemented in an OPM system, and our architecture facilitates light-sheet focusing over a depth range of 1 mm. The system’s performance is evaluated by imaging fluorescent nanospheres in agarose gels, and we demonstrate its application potential by imaging the vasculature and blood flow in zebrafish embryos. The spatial resolution of the system allowed us to resolve subcellular details such as endothelial nuclei and shape changes of red blood cells. Our quantitative blood flow measurements across a Zebrafish embryo indicated the need for acquisition rates above 10Hz. This further demonstrates the need for rapid mesoscopic imaging techniques.

## 2. Methods

### 2.1 Light-sheet reflection by a microprism

Reflection of a light-sheet has been previously accomplished with a mirror near the sample in techniques such as soSPIM (Figure 1A). To translate this concept to an open top, mesoscopic OPM system, the reflection of the light-sheet must occur below the sample. This can be accomplished with a knife edge mirror (Figure 1B) or via reflection in a higher refractive media (Figure 1C). When using the knife edge mirror with an air objective, the following constraints result due to Snell’s law: to achieve a desirable tilt angle of 45° of the light-sheet in water, a 20° incidence beam at the air-glass interface is required. This means that the light travels at a shallow angle to the coverslip and the presence of a small gap between the mirror and coverslip would lead to a considerable distance traveled laterally before entering the sample. This in turn limits the field of view over which the beam can be scanned. In addition, a 45° tilt angle would also be close to the maximum tilt angle (48.6°) that could be achieved in such a configuration, as a result of refraction.

**Fig. 1.**
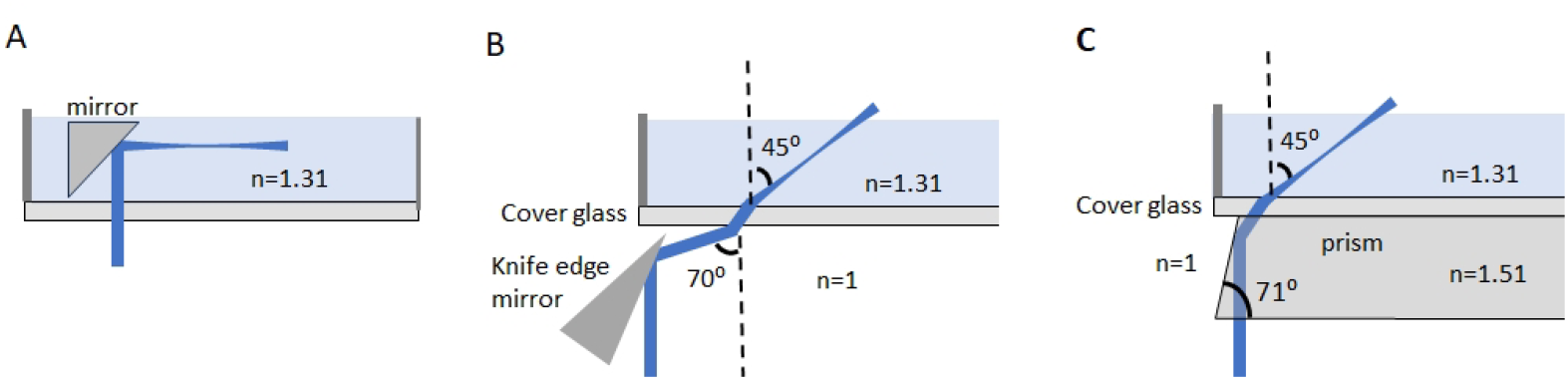
Light-sheet reflections in different implementations. **A** In soSPIM, a mirror reflects a light-sheet by 90 degrees. **B** Using a knife edge mirror to tilt a light-sheet in the sample space. **C** Using total internal reflection in a glass prism to tilt a light-sheet. **B-C** assume air objectives.

In contrast, in an optically denser medium such as a glass prism, the laser light travels at a steeper angle before entering the watery medium (Figure 1C), resulting in a larger field of view that can be accessed. Further, the light-sheet tilt angle can in principle reach up to 90□ in water when the total internal reflection condition is reached. Realistically, tilt angles of 60-75□ (to the coverslip normal) are possible, which might be of interest for mesoscopic OPM applied to shallow samples.

Importantly, the mirror reverses the scan direction and modifies the scan amplitude of the light-sheet (Supplementary Figure 1). As such, the light-sheet is scanned in the opposite direction than the fluorescent light is de-scanned. A second consequence of the light-sheet mirroring is that the beam waist will be axially shifted as it is laterally scanned, an effect that is also present in soSPIM [21] (Supplementary Figure 2 and Supplementary Note 1). Both effects are compensated for in our experimental setup as detailed below.

### 2.2 Optical setup

A schematic of the optical layout of our mesoscopic OPM system is shown in Figure 2. Laser light (blue) is shaped into a light-sheet by the illumination engine [23]. Two galvo mirrors (QS20Y-AG, Thorlabs) in an image space between the first and second tube lens (TL1 and TL2, Both ITL 200, Thorlabs) perform “lens-less” scanning of the laser beam. Compared to traditional galvo scanning in OPM, this arrangement dispenses with two scan lenses, which results in less complexity and higher light-throughput (a detailed analytical description is given in Supplementary Note 2). The light-sheet emerges from the primary objective (O1, Olympus XLFLUOR4X/340, 4X magnification, Numerical aperture 0.28) along the optical axis and undergoes total internal reflection in a microprism (Perkins Precision, BK 7, 4.2mm thickness, see also subpanel i in Figure 2). Placing a slab of glass in front of an objective would typically result in spherical aberrations. However, the used primary objective is designed to image into a 5 mm deep water column. As such, a glass plate of equivalent optical path length can be introduced without causing deleterious spherical aberrations. In fact, without such a tall water column or glass plate, spherical aberrations do occur [24]. For this reason, an additional optical flat is placed after O2, and the grating substrate compensates spherical aberrations for O3. The proper balancing of spherical aberrations was verified by imaging fluorescent beads with widefield based illumination; spherical aberrations are minimized when blur rings are symmetrical above and below the central body of the PSF.

**Fig. 2.**
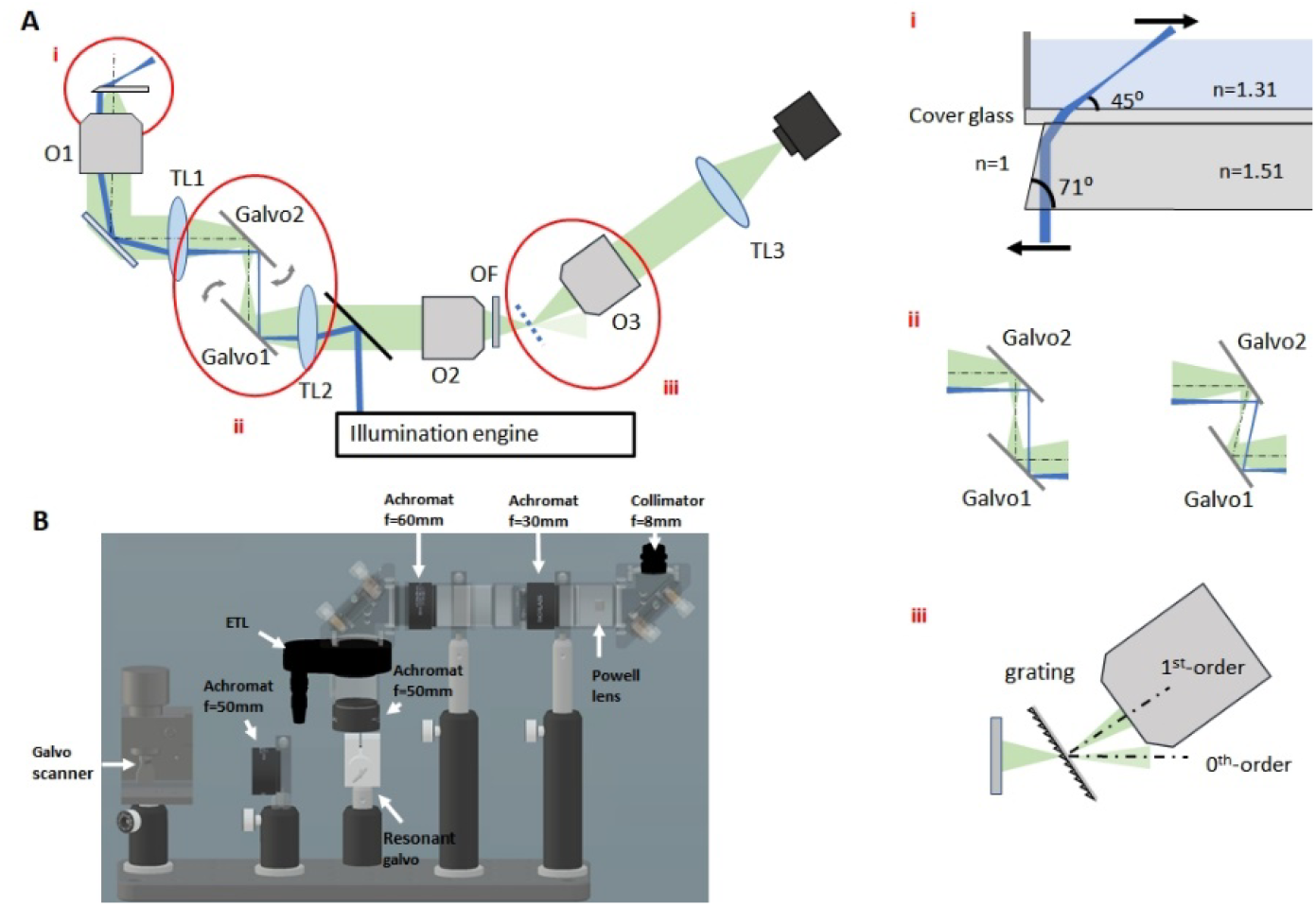
Schematic setup of the mesoscopic oblique plane microscope. **A** O1-O3: primary, secondary and tertiary objective. TL1-TL3: primary secondary and tertiary tube lens. OF: Optical Flat. Inset i shows detail of the microprism that reflects the light sheet into the sample. Inset ii shows the working principle of the image space scanning. Inset iii shows how the blazed diffraction grating diffracts the first order towards the primary objective. **B** Rendering of the illumination engine. The resonant galvo is located in a conjugate image plane, the Galvo scanner is conjugate to the pupil of the primary objective O1.

Fluorescent light captured by the primary objective is de-scanned by the galvo mirror pair and re-imaged with the secondary objective (O2, Olympus XLFLUOR4X/340, 4X magnification, Numerical aperture 0.28). A transmission grating (1200gr/mm, GT13-12, Thorlabs) diffracts the fluorescent light into the tertiary objective (O3, Olympus XLFLUOR4X/340, 4X magnification, Numerical aperture 0.28), which is then imaged via the tertiary tube lens (250mm focal length achromat, Thorlabs) onto an ORCA Flash 4 (2048×2048 Hamamatsu) or Kinetix (3200×3200 pixels, Photometrics) sCMOS camera. The overall magnification of the system is 5.65X, which ensures Nyquist sampling.

The illumination engine is shown in Figure 2B. A Powell lens (10□ fan angle, Laserline Optics, Canada) and a f=30mm achromatic lens (Thorlabs) are used to create a light-sheet. An electro tunable lens (ETL) and a galvo mirror, both conjugate to the back focal plane of the primary objective, are used to refocus and laterally scan and position the light-sheet, respectively. A resonant galvo, conjugate to an image plane, is used for shadow suppression [25]. The galvo mirror reverses the scan direction of the light-sheet and keeps it in lockstep with the fluorescence de-scan. An offset applied to the same galvo mirror is used to fine align the light-sheet position to the focal plane of the tertiary objective. The ETL dynamically compensates for the shift of the beam waist during lateral scanning. Further, the ETL can also axially shift the light-sheet on demand, which we leverage for tiling light-sheet microscopy. With focus compensation, we estimate the lateral scan range of 1.51mm using the 4.2mm thick microprism (Supplementary Note 1 and Supplementary Figure 2). Laser light is provided by a fiber coupled LightHUB Ultra light engine (Omicron Laserage Laserprodukte, Germany).

### 2.3 Transmission grating to diffract light into the tertiary objective

The use of diffraction gratings to guide light into the shallow acceptance cone of a tertiary objective was first shown by Hoffmann et al [16]. The idea is to place a grating at the desired image plane angle (i.e., along the image of the light-sheet) and choose the grating pitch such that (typically) the first order is normal to the grating surface. Using a reflection grating, a very steep light-sheet plane was picked up in the first demonstration of diffractive OPM. In our case, the image plane is less inclined, owing to the tilt angle augmentation of the light-sheet. This allowed us to use a transmission grating instead of a reflective grating and use a lower linespacing (1200gr/mm compared to 1800gr/mm). The grating compresses the light-cone in the diffraction direction [17], which yields to an elliptical light-distribution in the pupil of the tertiary objective. However, this effect is lessened for gratings of lower linespacing, and in our system leads to a fairly symmetric point spread function, even compared to conventional OPM systems that incur some levels of beam clipping [26].

The grating surface, where the diffraction happens, faces forward (Figure 2, inset iii). This was chosen because the grating is on a glass substrate, which has the potential to induce Coma aberrations when traversed at an angle. With the diffraction happening at the front surface of the substrate, the central ray travels close to orthogonally through the glass slide, thereby minimizing such aberrations. Experimentally, the diffraction efficiency into the first order remained ∼25% if the grating faced forward or backwards for a wavelength of 514 nm. Off note, the remote image space is slightly compressed in the third dimension, as we chose a unity lateral magnification from sample space to remote space, i.e., the primary and secondary lens, as well as the primary and secondary tube lens are the same (see also Supplementary Note 3 and Supplementary Figures 3-5).

## 3. Results

### 3.1 Imaging of fluorescent nanospheres

We first imaged 500 nm fluorescent beads (Invitrogen, FluoSpheres™ F8813) in 2% low melting agarose (Sigma Aldrich, A9045) in a placed in a Matek dish with a No 0 coverslip (Matek P35G-0-10-C). The dish was contacted to the microprism with a thin oil layer (n=1.52). We used such bead samples to calibrate the ETL based beam waist compensation and to assess the imaging performance of the system (Figure 3A-B).

**Fig. 3.**
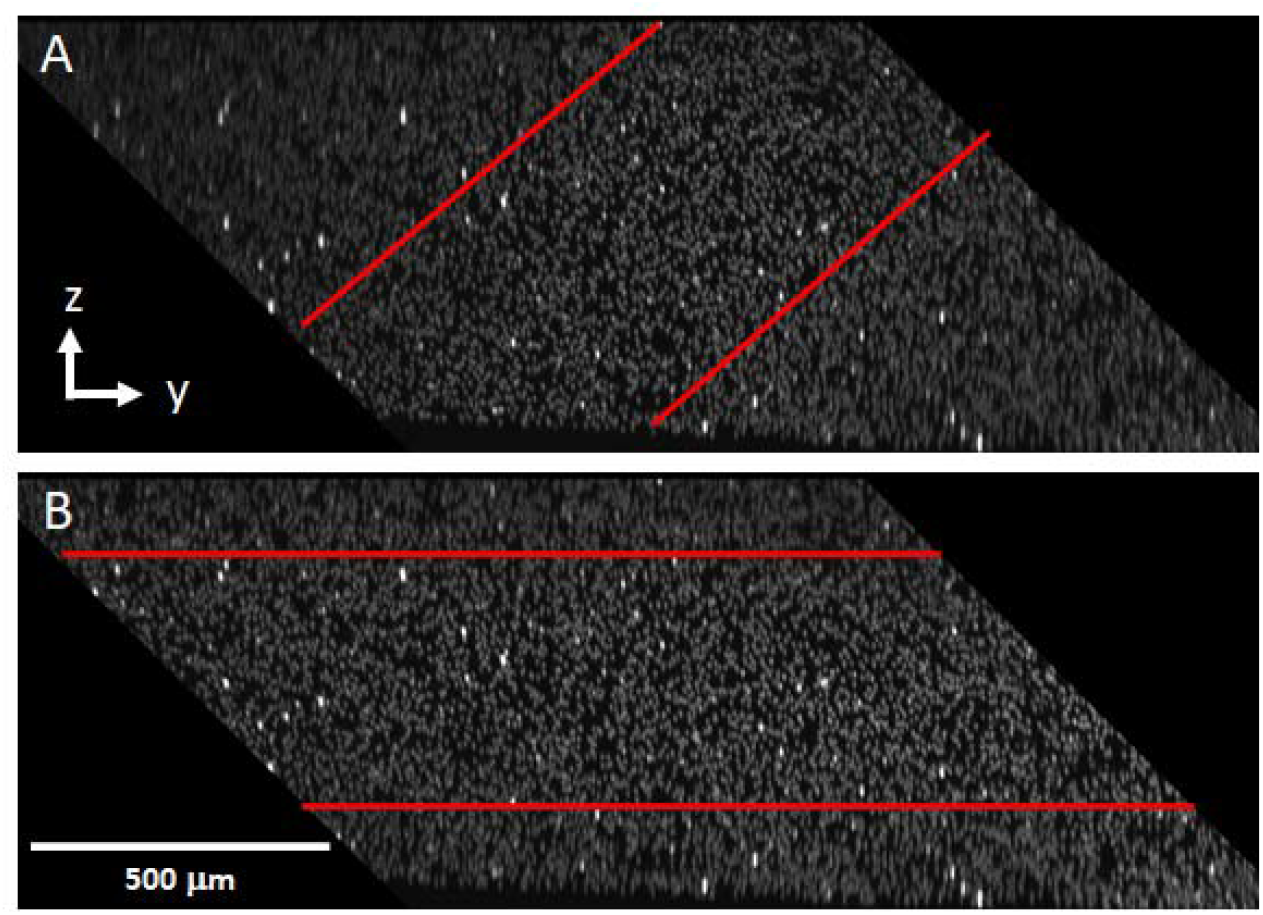
Beam waist position during stack acquisition, visualized by imaging 500nm fluorescent nanospheres in Agarose. **A** Without correction, beam waist is defocused during stack acquisition. **B** By refocusing with the ETL, the beam waist is held at a constant distance to the coverslip during a scan. Red lines in **A** and **B** help visualize the usable range of the light-sheet’s beam waist.

To estimate the resolution, we analyzed the full width half maximum (FWHM) of 500 nm fluorescent nanospheres in a 2% low melting agarose gel over a depth of 330 microns (which corresponds to the extend of the beam waist in the z-direction). Analysis of 1720 beads resulted in 2.33 ± 0.37μm, 2.77±0.32μm and 10.16±1.49μm for the FWHM in the x-, y-and z-direction, respectively (mean ± standard deviation). As shown in Figure 4A-B, the point-spread functions remained constant across the field of view.

**Fig. 4.**
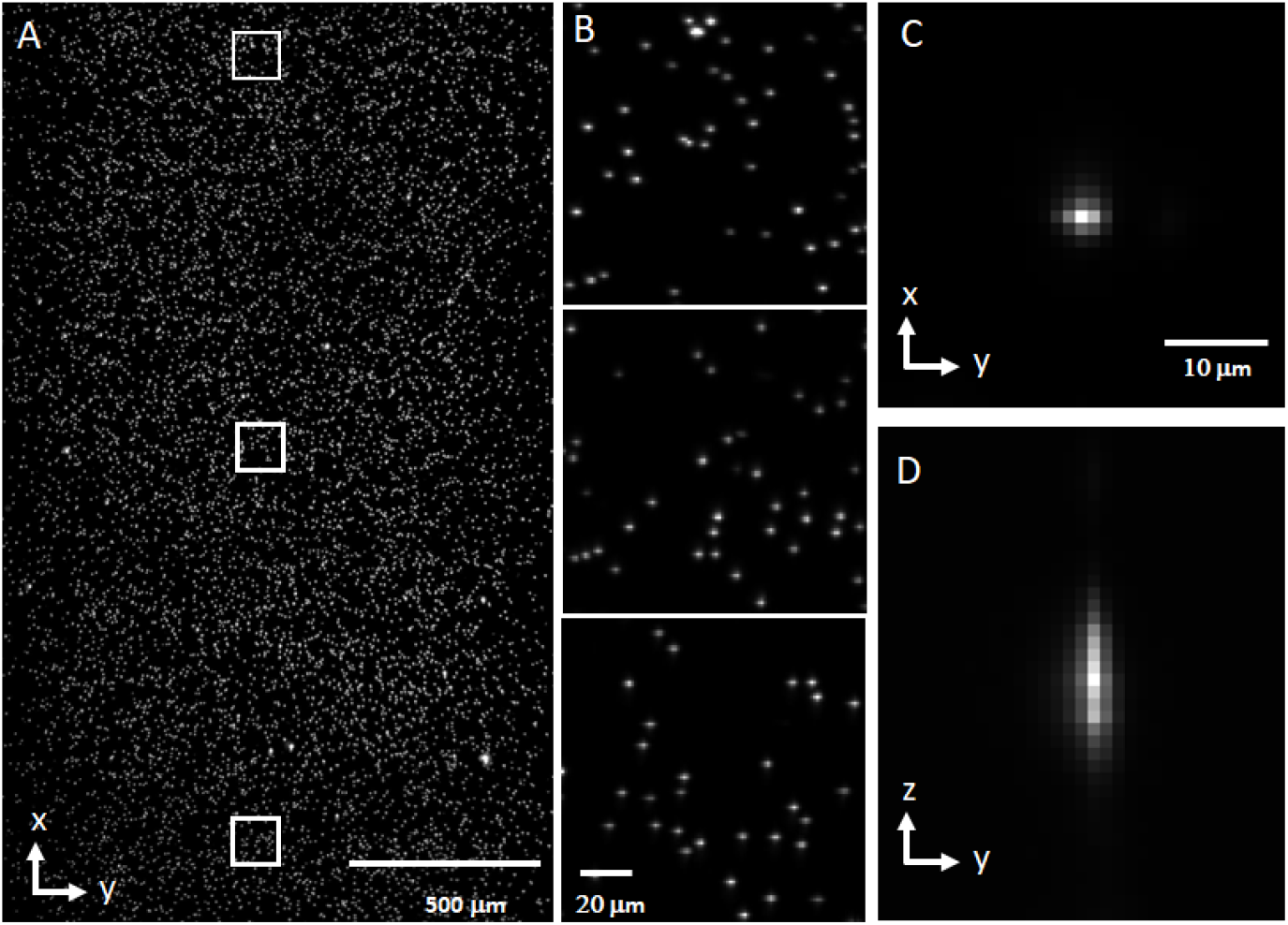
Imaging of 500nm fluorescent nanospheres in Agarose. **A** Maximum intensity projection over a 330-micron depth. **B** Magnified views of the white boxes in **A. C-D** Point spread function selected from one nanosphere image.

As with any light-sheet microscope employing Gaussian beams [27], axial resolution and length of the usable beam waist are coupled. In our system, we set the NA of the light-sheet to 0.04. Since the light-sheet propagates along a plane tilted by 45 degrees to the coverslip, the “height” above the coverslip (i.e., the imaging depth) that is covered with the beam waist is further reduced by a factor of ∼0.707. Increasing the depth coverage could be achieved by lowering the excitation NA, which would reduce the axial resolution. Alternatively, one can refocus the light-sheet waist to different depths for tiling, which has been implemented in conventional LSFM configurations [22], but not in OPM to our knowledge. One aspect that complicates the implementation in OPM is that the light-sheet is typically composed of marginal rays of a high NA objective, which would require complex wavefront shaping for proper re-focusing. In our case, the lights-sheet leaves the objective along its optical axis, and as such makes refocusing analogous to refocusing a light-sheet in a conventional LSFM architecture. This appeared to work well to dynamically stabilize the waist position during scanning using small refocus amounts, and hence we explored if we could use the same mechanism also for tiling microscopy. We imaged six individual stacks with different axial waist positions (Figure 5A) over a depth range of 1 mm, and computationally fused the volumes to a single stack (Figure 5B). The individual stacks were fused with a custom Python script using weighted average fusion based on a sigmoidal function. The axial resolution over a height of 1mm was maintained at 9.225 ± 1.151 μm (mean ± standard deviation, n=417).

**Fig. 5.**
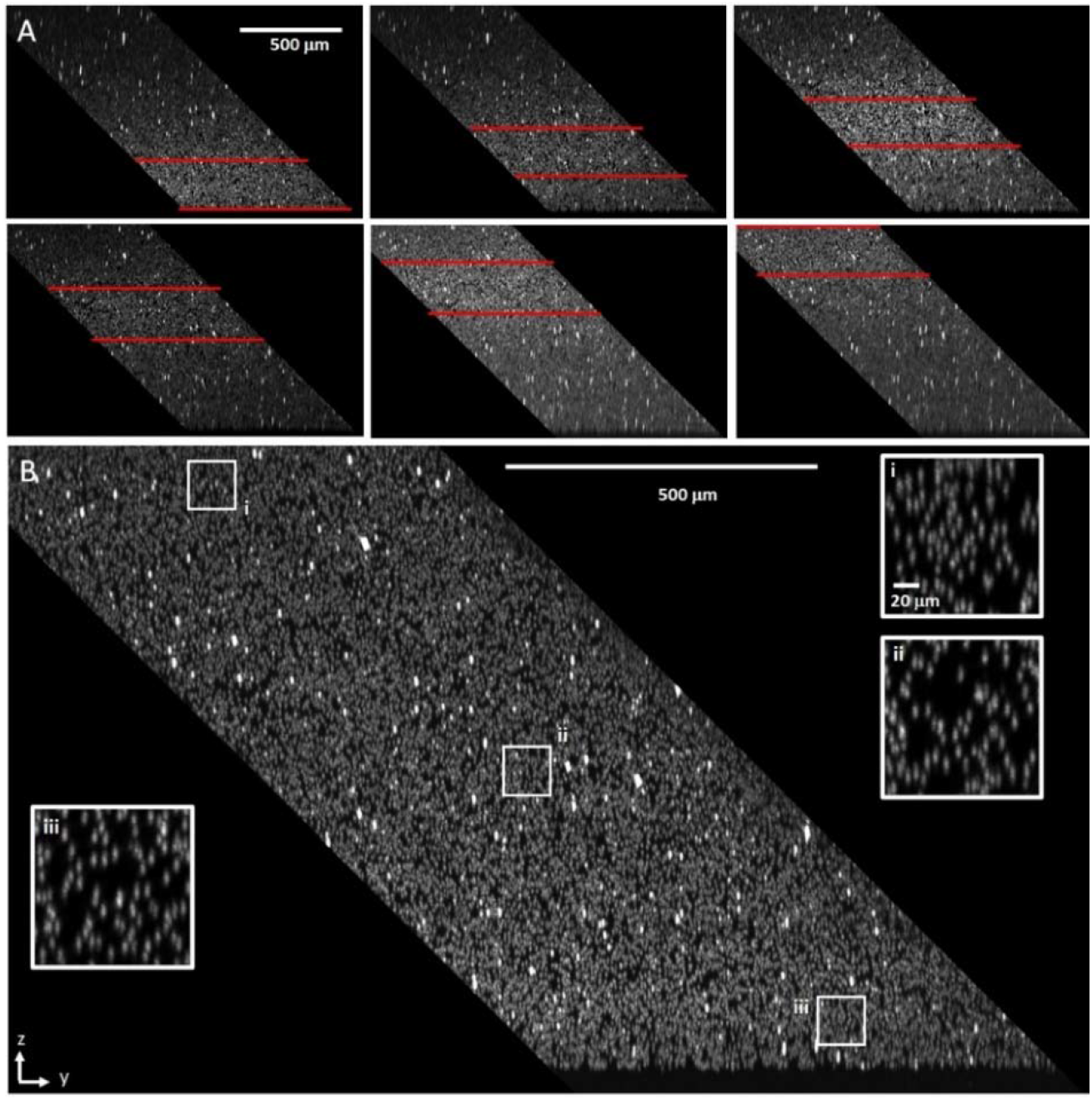
Increasing the depth range while maintaining constant z-resolution via optical tiling. **A** Six consecutive volumes are shown where the z-position of the beam waist was varied, starting at the bottom (near the coverslip). Red lines indicate the beam waist location. **B** Fusion of the six tiles. Insets show magnified views of the boxes near the bottom, middle and top of the fused volume.

### 3.2 Imaging Zebrafish embryos

To demonstrate the intravital imaging capabilities of our mesoscopic OPM system, we performed fast live imaging of entire zebrafish embryos at three days post fertilization. Zebrafish expressed the vascular marker Tg(*kdrl:EGFP*) and the red blood cell marker Tg(*gata1a:DsRed*) in a casper background [28]. To mount the zebrafish embryos, they were anesthetized with 200 mg/l Tricaine (Sigma Aldrich, E10521) and placed in a Matek dish with a No 0 coverslip (Matek P35G-0-10-C) at the bottom. A second glass coverslip on top provided further mechanical support during imaging. Zebrafish husbandry and experiments followed established protocols and have been approved and conducted under the oversight of the Institutional Animal Care and Use Committee (IACUC) at UT Southwestern under protocol number 101805 to Gaudenz Danuser.

The field of view of the mesoscopic OPM system covered the whole zebrafish embryo vasculature (Figure 6A-F). To obtain higher resolution and better depth coverage, we fused two tiled volumes. As with any LSFM technique, light-scattering by the sample induced some blur further away from the detection objective (Figure 6D). Nevertheless, the head and tail vasculature of the zebrafish embryo were well sectioned throughout the volume (Figure 6D-F). Importantly, the mesoscopic OPM system resolved subcellular details (magnified views in Figure 6B-C and E-F), including nuclei of endothelial cells (bright, localized spots) and the intricate branching patterns of zebrafish vasculature, including parachordal lymphiangioblasts.

**Fig. 6.**
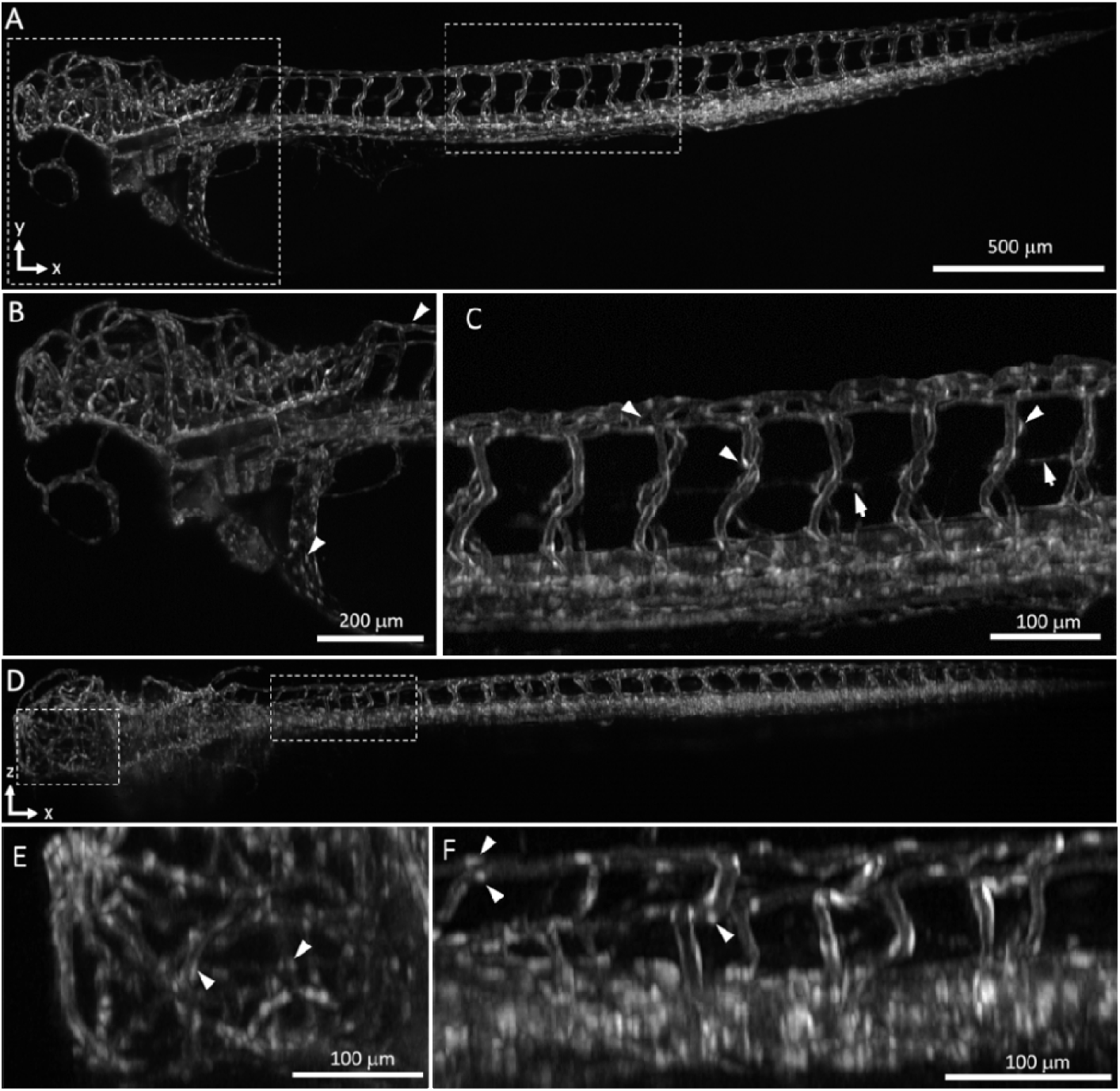
Imaging of Zebrafish vasculature. Fluorescently labeled vasculature, *Tg(kdrl:EGFP)*, in a three days post fertilization (dpf) zebrafish embryo, as imaged with our mesoscopic OPM. **A** X-Y maximum intensity projection of the entire zebrafish with **B-C** Magnified views (head and tail vasculature) of the boxed regions in **A**. Arrowheads indicate selected endothelial nuclei and arrows point to parachordal lymphangioblasts. **D** X-Z maximum intensity with **E-F** X-Z maximum intensity projected magnified views (head and tail vasculature) of the boxed regions in **D**. Arrowheads indicate selected endothelial nuclei.

To demonstrate the potential for rapid volumetric imaging, we imaged blood flow dynamics without optical tiling (one stack was acquired per timepoint). With 4×4 binning, we could achieve a volumetric imaging rate of 5Hz. While it was possible to identify and track blood cells manually with anatomical knowledge of zebrafish vasculature, 5 Hz was too slow for automated analysis of blood flow dynamics with optical flow analysis (Supplementary Note 4). Therefore, we leveraged our recently introduced projection imaging technique [29], where the light-sheet is rapidly swept through the 3D volume, and the fluorescence image is synchronously scanned (‘sheared’) over the image sensor. To this end, we added a single galvanometric mirror (Thorlabs QS20Y-AG) in front of the camera for optical shearing. Together, this enabled rapid time-lapse projection imaging at 12 Hz imaging rate with coverage of the whole embryo over one hundred timepoints.

The resulting imaging data enabled qualitative and quantitative analysis of blood flow across the entire embryo (Figure 7). As blood cells are sparse objects in the blood stream, we applied maximum intensity projections over the time-lapse to obtain a more continuous map of vessel perfusion (Figure 7A, Supplementary Movie 1). Moreover, color-coding of subsequent time-points (Figure 7B) revealed the directionality of flow within the vessels. Thereby, the mesoscopic OPM system also captured subcellular details such as subtle differences of red blood cell shapes in different vessels (Figure 7C-F). To quantitatively determine the blood flow direction and magnitude across a whole embryo, we computed multiscale optical flow using Farnebäck’s method [30], implemented in OpenCV [31] to measure the frame-to-frame blood cell movement. As movement is only measured in presence of cells, we sampled the velocity vector corresponding to the 95th percentile speed measured over the video duration to reconstruct the underlying blood flow field at each pixel position (Figure 7G, Supplementary Note 4). Our analysis clearly resolved the directionality of blood flow within the whole zebrafish vasculature present in the projection images.

**Fig. 7.**
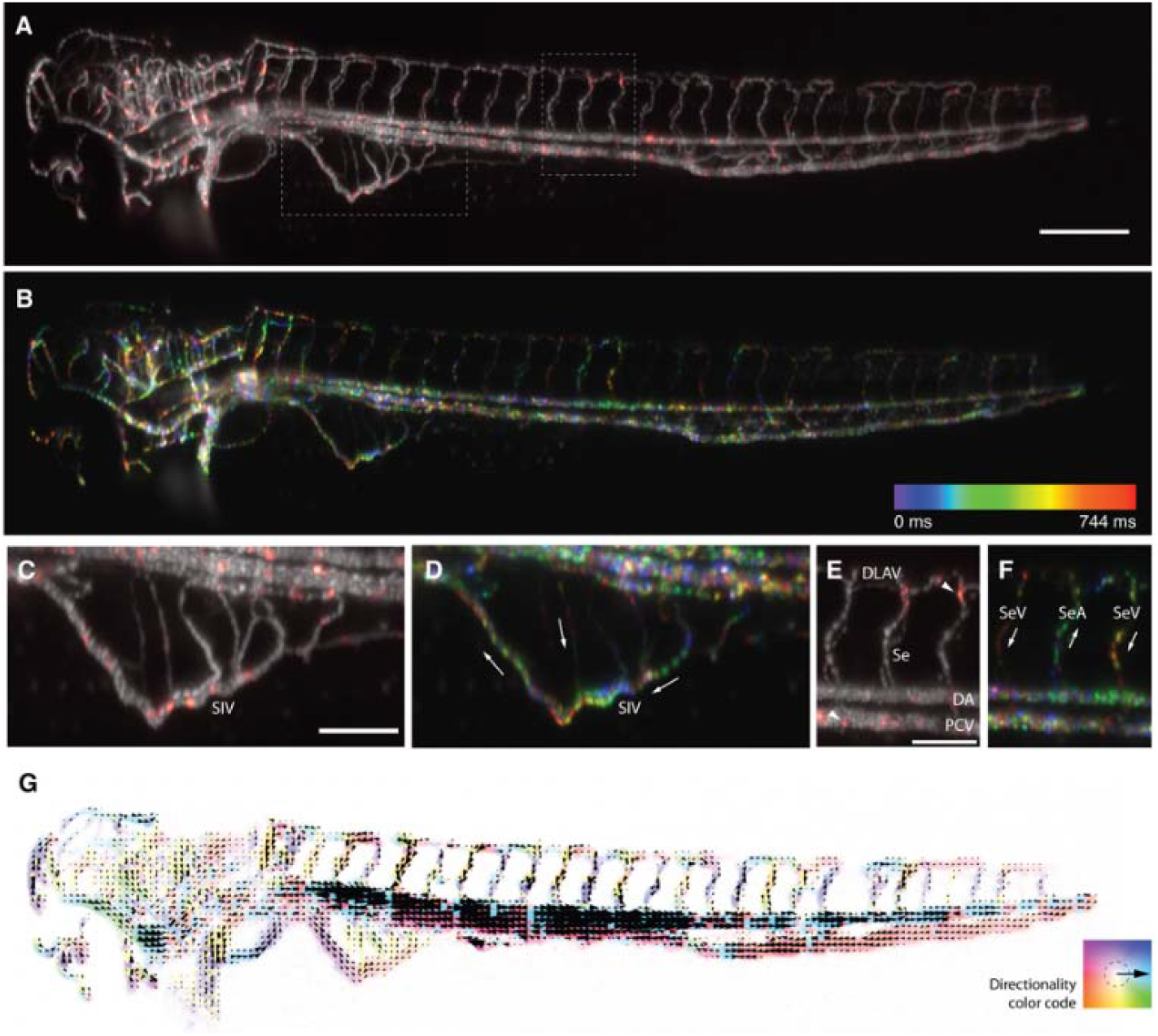
Imaging and quantification of blood flow in zebrafish embryos. Blood flow dynamics in a 3 days post fertilization (dpf) zebrafish embryo, as imaged with our mesoscopic OPM in a projection format at 12Hz over 100 timepoints. **A** The maximum intensity of the hundred frames (gray) provides a visual impression of the vasculature and a map of which vessels were perfused. In red, a single time point of the movie is shown from which the average signal over the time-lapse was subtracted to highlight individual, bright red blood cells. Insets depict the regions which were magnified in C-F. Scale bar: 250 μm. **B** Ten subsequent timepoints of the movie (744 ms) were color-coded and overlaid on each other **C**,**D** Magnified views of the subintestinal vein (SIV) plexus. White arrows indicate blood flow direction. Scale bar: 100 um. **E, F** Magnified views of the dorsal longitudinal anastomotic vessel (DLAV), three intersegmental vessels (Se), the dorsal aorta (DA), posterior cardinal vein (PCV). Scale bar: 100 μm. **E** White arrowheads highlight that red blood cells adopt different shapes in different vessels, including elongated shapes in intersegmental vessels and compact, spherical shapes in large vessels with fast flow. **F** Color coding further allowed to identify the vessel identity and distinguish intersegmental artery (SeA) and intersegmental vein (SeV). White arrows indicate blood flow direction. **G** Result of quantitative analysis of blood flow with optic flow using Farnebäck’s method. The color code and arrow orientation indicates the directionality of the flow, and arrow length and color saturation the magnitude of the flow.

## 4. Discussion

We have introduced a mesoscopic OPM system that combines concepts of soSPIM, diffractive OPM and tiling light-sheet microscopy, as well as a new lens-less galvo scanning mechanism. Taken together, this resulted in a compact, easily aligned system that improves 3D imaging performance over previous mesoscopic OPM systems. To demonstrate the power of our mesoscopic OPM system, we imaged embryonic zebrafish vasculature and blood flow dynamics *in vivo*, resulting in a quantitative map of blood flow dynamics across an entire embryo.

A key element in our mesoscopic OPM design is the microprism, which augments the light-sheet tilt angle via total internal reflection. Thereby, we can create light-sheet tilt angles which exceed the half opening angle of the primary objective. We chose a light-sheet tilt angle of 45 degrees, as it forms a good compromise between optical sectioning and volumetric coverage (i.e., the height above the coverslip that can be covered with a given beam waist). Nevertheless, our method can also allow higher tilt angles. As an example, a prism angle of 61 degrees instead of 71 degrees would result in a light-sheet that is tilted by ∼ 30 degrees to the coverslip.

Importantly, our approach via total internal reflection is wavelength independent. This is a considerable advantage compared to using a diffraction grating close to the sample plane, as introduced by Shao et al [19], which induces a wavelength dependent tilt angle. Consequently, we can perform multicolor imaging without modifying the setup, in contrast to Shao et al who needed to change the tilt of the tertiary imaging system for different colors. The grating will also create unwanted higher diffraction orders with the fluorescent light. While they do not appear prominently in the images presented by Shao et al, they still subtract valuable signal from the zero order. Lastly, a zero-diffraction order of the excitation light needed to be blocked with a moving mask, which will limit the ultimate speed of the grating method. In contrast, total internal reflection occurs with such a high efficiency that we did not need to block any unwanted excitation light.

Besides the aspects of efficiency and wavelength dependency, there are geometrical effects. Light-sheet scanning can occur anywhere across the grating. In contrast, in our method, light-sheet scanning must occur across the tilted face of the prism, so more care for adjusting the position must be taken as the tilted flank of the prism cannot be used for fluorescence imaging. Additionally, the total internal reflection on the prism causes a reversal of the scan direction, and a defocusing of the light-sheet. In our setup, the reversal of the scan direction is countered with the light-sheet positioning mirror in our illuminator. This mirror has so far always been motorized in our setups, as it is used to align the light-sheet position to the focal plane. Thus here, the mirror not only ensures the proper alignment of the two but is also used for compensating for the scan reversal. Of note, it is also conceivable to place the dichroic right after the primary objective and use this space to couple in the light-sheet. Our solution minimized in comparison the total number of lenses needed.

Furthermore, the electro tunable lens (ETL) in our illuminator introduces optical tiling to OPM systems besides its function to compensate the defocusing of the light-sheet waist. As demonstrated here, this can extend the reach of our mesoscopic OPM in the third dimension up to 1mm, about three times the range shown by Shao et al. We believe that this capability is of importance for large transparent organisms, or samples that are not attached to the coverslip surface, which may include freely swimming embryos. Lastly, expansion microscopy and tissue clearing now produce very large transparent samples. As such, improvements of the axial reach are important for the next generation of mesoscopic OPM systems.

The lateral width of the field of view is currently limited by the camera to enable Nyquist sampling. For a 3200×3200 pixel camera, a width of ∼3,680 mm results. With cameras with more pixels, a 5mm width might be possible given the field of view of the objectives used. In the scan direction, the prism imposed a limit of 1.51 mm. The glass prism thickness could be doubled if the optical flat after the secondary objective were left away. This can be seen as follows: somewhere in the system, an aberration correction for a combined 15mm deep water column must occur. This correction can occur at any stage, i.e. in front of the primary objective, in the remote space, or in both spaces. Assuming a FOV of 5 mm of the primary objective, this would enable ∼2.78mm scan range (2.21mm would be used up by the inclined flange of the prism).

The high sampling requirements may slow down acquisition speed. We have countered this by pixel binning and projection imaging. While these methods make compromises in terms of spatial sampling and dimensionality of the data, they can potentially be advantageously applied in a multi-modal fashion [32]. As an example, one camera channel could record the entire fish at the full 3D resolution, while another channel either covers a sub volume (like the brain) or features that require less resolution (like blood cells), or processes that can be interrogated by 2D projection imaging.

In conclusion, we have introduced a compact mesoscopic imaging system that features large volumetric coverage and comparatively high spatial resolution. This has been enabled by light-sheet mirroring to increase its tilt angle, the use of a transmission grating to diffract light into the acceptance cone of the tertiary objective, and optical tiling. An image-based scanning mechanism further simplifies the optical train of our system. As such, we hope that this system will find widespread applications in biological and biomedical research, as it is easy to build and offers high spatiotemporal resolution in the mesoscopic imaging realm.

## Acknowledgments

The authors are thankful to Dr. Fabian Voigt for fruitful discussions. The authors are also grateful to the Danuser lab at UT Southwestern for their help on zebrafish husbandry and computational support. Moreover, this research was supported in part by the computational resources provided by the BioHPC supercomputing facility located at UT Southwestern Medical Center.

## Data availability

Data underlying the results presented in this paper is publicly shared on this Zenodo repository: https://zenodo.org/record/8200953

## Supplemental document

See Supplement 1 for supporting content.

## SUPPLEMENTAL DOCUMENT

### Supplementary Notes

#### Supplementary Note 1 Analysis and simulation for light-sheet scan range

We analytically derived the relationship between scanning at the bottom of the prism and the resulting scanning at the top of the prism (Supplementary Figure 1).

Next, we used Zemax to analyze the illumination light sheet after the microprism. Without compensation of the ETL, the focus of the light sheet shifts along the scanning direction (Supplementary Figure 2A). This causes a tilted focal plane, also apparent in the acquired beads (Figure 3A). By adding an ETL, we compensated for the axial shift and tuned the focal plane parallel to the coverslip plane (Supplementary Figure 2B, bead data Figure 3B).

As shown in Supplementary Figure 2, the blue beam represents the light sheet along the main optical axis. The maximum scan range without beam clipping is -0.53 mm (green beam) to 0.66 mm (red beam) before the microprism (Δx in Supplementary Figure 1), and 0.67 mm to -0.84 mm after the microprism (Δs in Supplementary Figure 1). The non-symmetricity is caused by the divergence of the beam. The optical power of ETL used to compensate for the shift ranged from -0.135 diopter (dpt) to 0.169 dpt.

#### Supplementary Note 2 Dual galvo image scanning

Adding a pair of Galvo mirrors in front of a camera to shear the image has been demonstrated in [1] and is discussed in more detail in [2]. Here, we extended its application to realize lens-less sample scanning in oblique plane microscopy.

According to our Zemax simulation (Supplementary Note 1), the maximal scanning range that the prism slab can afford without beam clipping is from -0.53 to 0.66 mm after the primary objective, which corresponds to a scan range of 1.51 mm in sample space. In the image space, this corresponds to a lateral shift (x) of -2.36 to 2.93 mm for the dual galvo scanner (magnified by 200/45). The distance (d) between the galvo pairs is 60 mm. From [2], we have the mechanical scanning angle α of the dual galvo as

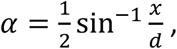

then the scanning angle is -1.13 to 1.4 degrees. The path length difference Δ compared to the center beam is

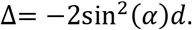

After demagnification of the primary objective, the defocus of the scanning edge is -3.13 and -4.83 um. We consider this effect to be negligible compared with the overall scan length of ∼1.51 mm.

#### Supplementary Note 3 Compression of the remote image space

In our OPM system, the lateral magnification from sample to remote space equals to 1. The remote space is in air, and has a refractive index of 1, whereas the sample is aqueous. As such, the axial magnification of the remote image equals to[3]:

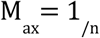

With n denoting the refractive index of the sample. Supplementary Figure 3 depicts the axial compression of a square sample volume.

Due to the compression, angles are not maintained. The change of the light-sheet angle α?in sample space to the remote space α’ can be expressed as:

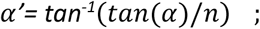

With α’ being the angle in the remote space, see also Supplementary Figure 4. For a light-sheet angle α =45° and n=1.33, the light-sheet angle α’ in remote space is reduced to ∼36.9°.

Besides the reduction of the light-sheet tilt angle, the light-sheet plane that is imaged by the tertiary objective is also compressed in one dimension (**Supplementary Figure 5)**. The compression factor equals to:

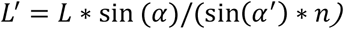

For a=45° and n=1.333, we find L’∼0.884*L.

#### Supplementary Note 4 –required imaging speed for blood flow analysis

To quantitatively determine the blood flow directionality and magnitude across the entire embryo given the imaged blood cells, we used multiscale optical flow computed using Farnebäck’s method [4]Thereby, we chose the following settings of the Farnebäck’s OpenCV [5] implementation: a classical Gaussian pyramid with each layer twice smaller than the previous one (pyr_scale=0.5), 5 pyramid layers (levels=5), an averaging window size of 5 (winsize=5), 5 iterations (iterations=5), a 5 pixel neighborhood to find the polynomial expansion in each pixel (poly_n=5), and 1.2 the standard deviation of the Gaussian that is used to smooth derivatives used as the basis of the polynomial expansion (poly_sigma=1.2).

To separate the exceedingly fast blood flow in two near-touching vessels in the 2D projections, the dorsal aorta (DA) and the principle cardinal vein (PCV), we first established a binary mask, separating the calculation of optical flow into two time-lapse series of the ventral and the dorsal parts, separated between DA and PCV. Subsequent stitching of optical flow vectors reconstructed the full optical flow of the whole embryo. Optical flow was computed as described and with the above parameters between every subsequent timepoint. As blood cells are spherical, sparse objects within the vasculature, blood cells are not always present in every frame at the same pixel location. In order to reconstruct the final continous estimate of blood flow velocity, we sampled the velocity vector corresponding to the 95^th^ percentile speed for every pixel over time.

To simulate the minimal necessary acquisition speed for automated blood flow estimation using optical flow analysis, we subsampled the acquired time-lapse of 100 frames, imaged at 12 Hz. Thereby, we retained every image (12.1 Hz), every second (6.1 Hz), every third (4 Hz), every fourth (3 Hz) and every fifth image (2.42 Hz) for comparison. By comparing the resulting image, we realized that 12 Hz was required as already at 6 Hz, the mean Euclidean distance (magnitude and direction; Supplementary Figure 6A) and Cosine distance (direction;

Supplementary Figure 6B) compared to the reference of 12.1 Hz increased considerably. Moreover, when visualizing the directionality of the flow at the different simulated acquisition speeds, differences were apparent and there are clear discrepancies with known zebrafish blood flow (Supplementary Figure 6C,D).

## Supplementary Figures

**Supplementary Fig. 1.**
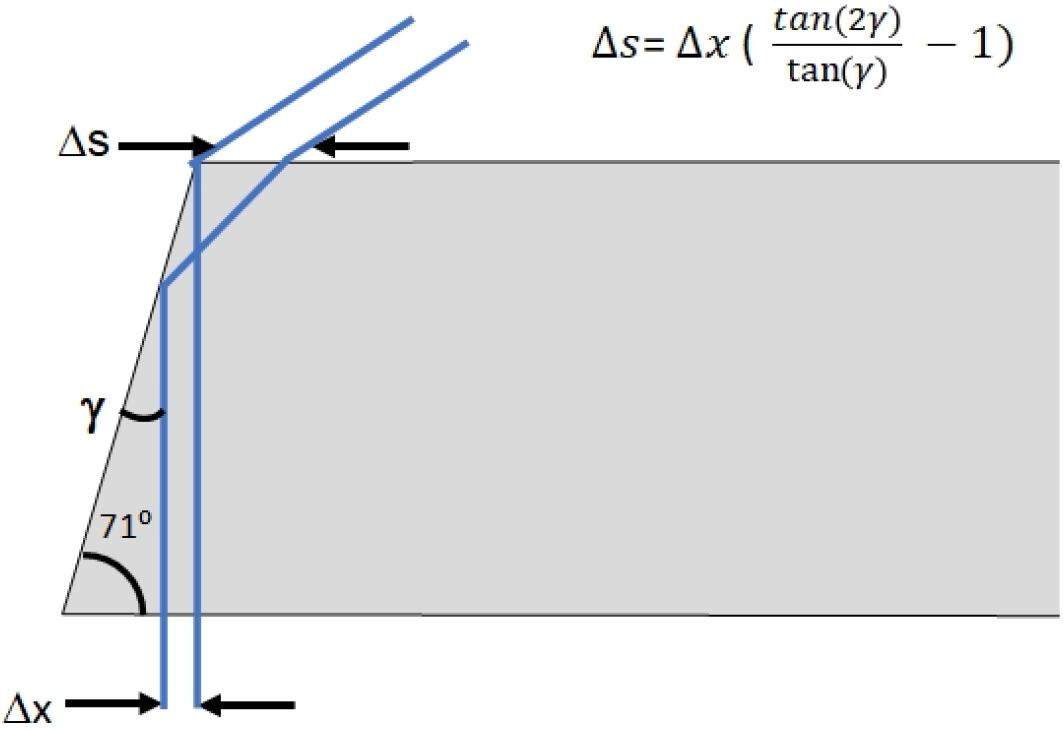
Relation between scanning at the bottom of the prism, and the resulting scanning at the top of the prism.

**Supplementary Fig. 2.**
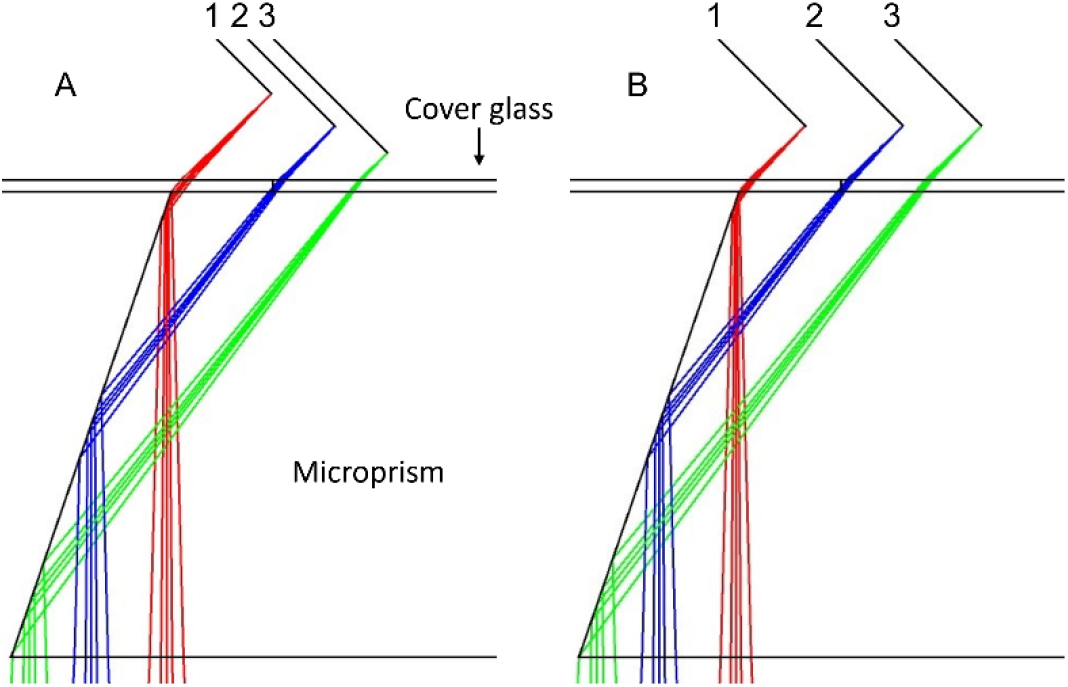
Zemax simulation of the microprism. **A** Beam waist shifts in axial dimension during the stack acquisition. **B** Beam waist shifts compensated with ETL. Colors present three configurations: beam along the main optical axis (blue / configuration 2) and scanned to both end without beam clipping (green / 3 and red / 1).

**Supplementary Fig. 3.**
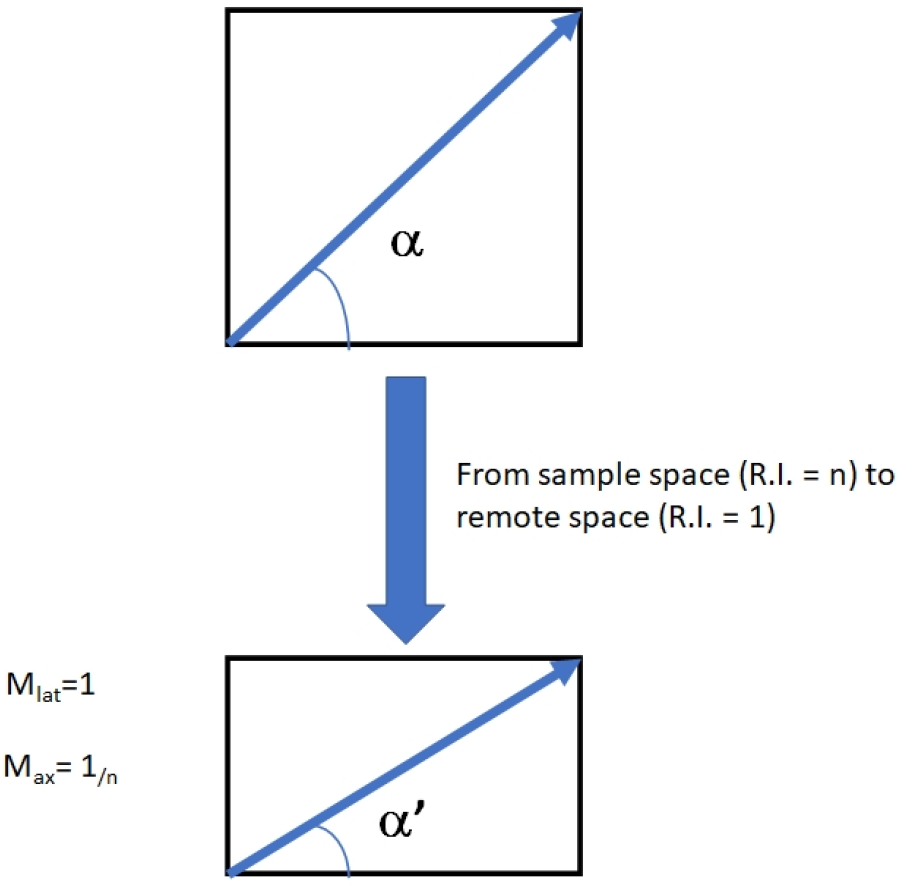
Compression of the sample space into the remote space when a lateral magnification of 1 is chosen and the refractive index (R.I.) in the remote space is 1, and the sample R.I. n>1. Blue line indicates the light-sheet plane.

**Supplementary Fig. 4.**
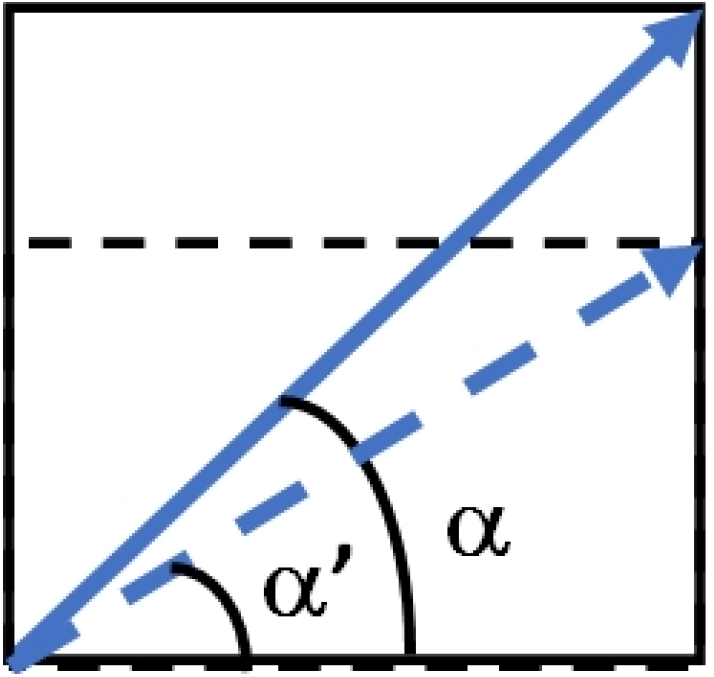
Compression of the light-sheet angle α to the remote space (dotted line, α’).

**Supplementary Fig. 5.**
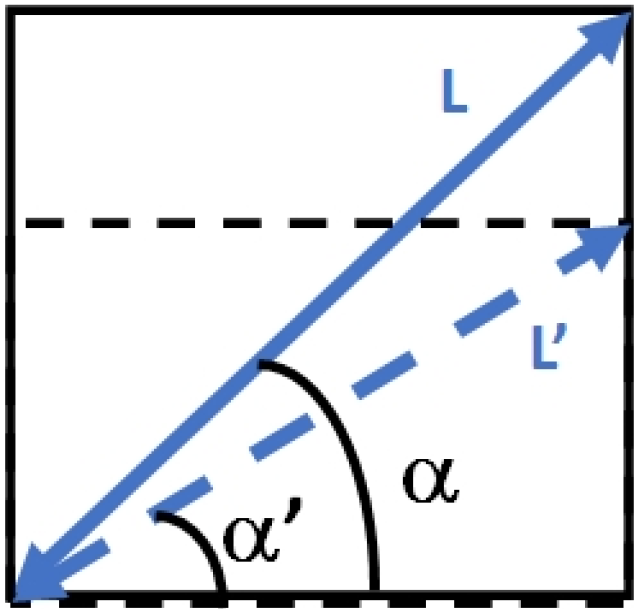
Compression of the L-axis from sample space to the remote space (L’). The L-axis runs along the propagation direction of the light-sheet.

**Supplementary Fig. 6.**
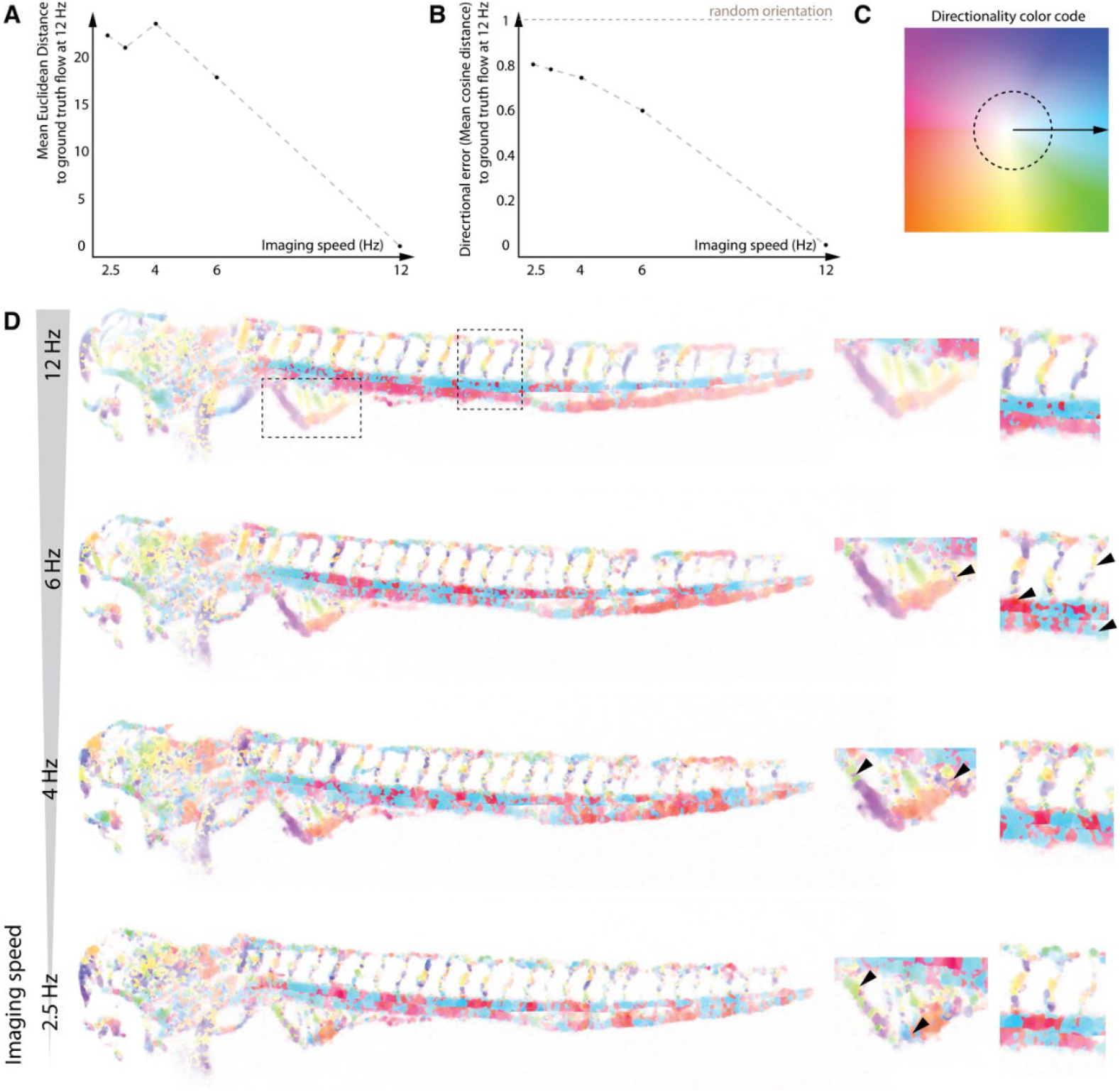
Analysis of the minimal imaging speed for automatic blood flow measurement using optical flow. **A** Plot of the mean Euclidean distance between the optical flow inferred blood flow at different simulated imaging rates relative to the reference blood flow determined at 12.1 Hz, restricted to the area of the segmented vasculature. **B** Plot of the directional error calculated as the mean cosine distance between the optical flow inferred blood flow at different simulated imaging rates compared with the reference blood flow obtained at 12.1 Hz imaging rate, restricted to the area of the segmented vasculature. **C** Directional color code map for panel D. The stronger the saturation, the faster the flow. **D** Directional color code map of the blood flow computed by optical flow at different simulated imaging rates. The magnified insets (boxed regions in the embryo at 12 Hz) highlight the subintestinal (SIV) plexus and three selected intersegmental vessels. With decreasing imaging rate, errors in the calculation of directionality accumulate (black arrowheads). While the slower blood flow in the subintestinal plexus permits almost error free calculations still at 6 Hz, the fast flow in the dorsal aorta and principal cardinal veins and intersegmental vessels show abundant errors. At 4 Hz, the optic flow also in the SIV plexus starts to show errors (black arrowheads).

